# Sensitivity of Quantitative Traits to Mutational Effects, Number of Loci, and Population History

**DOI:** 10.1101/008540

**Authors:** Joshua G. Schraiber, Michael J. Landis

## Abstract

When models of quantitative genetic variation are built from population genetic first principles, several assumptions are often made. One of the most important assumptions is that traits are controlled by many genes of small effect. This leads to a prediction of a Gaussian trait distribution in the population, via the Central Limit Theorem. Since these biological assumptions are often unknown or untrue, we characterized how finite numbers of loci or large mutational effects can impact the sampling distribution of a quantitative trait. To do so, we developed a neutral coalescent-based framework, allowing us to experiment freely with the number of loci and the underlying mutational model. Through both analytical theory and simulation we found the normality assumption was highly sensitive to the details of the mutational process, with the greatest discrepancies arising when the number of loci was small or the mutational kernel was heavy-tailed. In particular, fat-tailed mutational kernels result in multimodal sampling distributions for any number of loci. An empirical analysis of 7079 expressed genes in 49 *Neurospora crassa* strains identified 116 genes with non-normal sampling distributions. Several genes showed evidence of multimodality and/or skewness, suggesting the importance of their genetic architecture. Since selection models and robust neutral models may produce qualitatively similar sampling distributions, we advise extra caution should be taken when interpreting model-based results for poorly understood systems of quantitative traits.

## 1. Introduction

Questions about the distribution of traits that vary continuously in populations were critical in motivating early evolutionary biologists. The earliest studies of quantitative trait variation relied on phenomenological models, because the underlying nature of heritable variation was not yet well understood (Galton, 1883, 1889; Pearson, 1894, 1895). Despite the rediscovery of the work of Mendel (1866), researchers studying continuous variation in natural populations were initially skeptical that the Mendel’s laws could explain what they observed (Weldon, 1902; Pearson, 1904). These views were reconciled when Fisher (1918) showed that the observations of correlation and variation between phenotypes in natural populations could be explained by a model in which many genes made small contributions to the phenotype of an individual.

The insights of Fisher (1918) made it possible to build models of quantitative trait evolution from population genetic first principles. Early work focused primarily on the interplay between mutation and natural selection in the maintenance of quantitative genetic variation in natural populations, while typically ignoring the effects of genetic drift (Fisher, 1930; Haldane, 1954; Latter, 1960; Kimura, 1965).

However, genetic drift plays an important role in shaping variation in natural populations. While earlier work assumed that a finite number of alleles control quantitative genetic variation (e.g. Latter (1970)), Lande (1976) used the continuum-of-alleles model proposed by Kimura (1965) to model the impact of genetic drift on differentiation within and between populations. A key assumption of Lande’s models is that the additive genetic variance in a trait is constant over time. In fact, in finite populations the genetic variance itself is random; at equilibrium, there are still stochastic fluctuations around the deterministic value assumed by Lande, even if none of the underlying genetic architecture changes (Bürger and Lande, 1994).

Several later papers explored more detailed models to understand how genetic variance changes through time due to the joint effects of mutation and drift (e.g. Chakraborty and Nei (1982)). Lynch and Hill (1986) undertook an extremely thorough analysis of the evolution of neutral quantitative traits. They analyzed the moments (e.g. mean and variance) of trait distributions that arise due to mutation and genetic drift and provided several quantities that can be used to interpret variation within and between species and analyze mutation accumulation experiments.

Much of this earlier work has made several simplifying assumptions about the distribution of mutational effects and the genetic architecture of the traits in question. For instance, Lynch and Hill (1986), despite analyzing quite general models of dominance and epistasis, ignored the impact of heavy tailed or skewed mutational effects. While, in many cases, such properties of the mutational effect distribution are not expected to have an impact if a large number of genes determine the phenotype in question, it is unknown what impact they may have when only a small number of genes determine the genetic architecture of the trait. Moreover, when mutational effects display “power-law” or “fat-tailed” behavior, the impact of the details of the mutational effects may persist even in the so-called infinitesimal limit of a large number of loci with small effects. Finally, mutation accumulation experiments have produced skewed and/or leptokurtic samples of quantitative traits (Mackay et al., 1992), which is a direct motivation to relax assumptions on the mutational effects distribution.

Such deviations that stem from the violations of common modeling assumptions have the potential to influence our understanding of variation in natural populations. For instance, leptokurtic trait distributions may be a signal of some kind of diversifying selection (Kopp and Hermisson, 2006) but are also possible under neutrality when the number of loci governing a trait is small. Similarly, multimodal trait distributions may reflect some kind of underlying selective process (Doebeli et al., 2007) but may also be due to rare mutations of large effect.

We have two main goals in this work. Primarily, we want to assess the impact of violations of common assumptions on properties of the sampling distribution of a quantitative trait (e.g. variance, kurtosis, modality). Secondly, we believe that the formalism that we present here can be useful in a variety of situations in quantitative trait evolution, particularly in the development of robust null models for detecting selection at microevolutionary time scales. To this end, we introduce a novel framework for computing sampling distributions of quantitative traits. Our framework builds upon the coalescent approach of Whitlock (1999), but allows us to recover the full sampling distribution, instead of merely its moments.

First, we outline the biological model and explain how we can compute quantities of interest using a formalism based on characteristic functions. We then use this approach to compute the sample central moments. While much previous work focuses on only the first two central moments (mean and variance), we are able to compute arbitrarily high central moments, which are related to properties such as skewness and kurtosis. By doing so, we are able to determine the regime in which the details of the mutational effect distribution are visible in a sample from a natural population. Additionally, we explore the convergence to the infinitesimal limit and find that when “fat-tailed” effects are present, traditional theory based on the assumption of normality can lead to misleading predictions about phenotypic variation. Finally, to assess the impact of genetic architecture in natural populations, we identified for non-normal sampling distributions of gene expression among 49 *Neurospora crassa* individuals.

## 2. Model

The mechanistic model we construct has three major components: a coalescent process, a genetic mutational process that acts upon the controlling quantitative trait loci, and a mutational kernel that samples quantitative trait effect sizes. Together these processes generate the quantitative traits sampled from the study population while explicitly modeling their shared genetic ancestry. Although we opt for simple model components during this exposition, the model generally supports more realistic and complex extensions, such as population structure and epistasis.

We assume that we sample *n* haploid individuals from a randomly mating population of size *N*. Initially, consider a trait governed by a single locus and we will later extend the theory to traits governed by multiple loci. Let *µ* be the mutation rate per generation at the locus, and *θ* = 2*Nµ* be the coalescent-scaled mutation rate. We model mutation as a process by which a new mutant adds an independent and identically distributed random effect to the ancestral state. Note that when the distribution of random effects is continuous, this corresponds to the Kimura (1965) continuum of alleles model. However, it is also possible for the effect distribution to be discrete, similar to the discrete model of Chakraborty and Nei (1982). While this model does not capture the impact of a biallelic locus with exactly two effects, the following theory could easily be modified to analyze that case.

Figure 1 shows one realization of both the coalescent and mutational processes for a sample of size 5. Given the phenotype at the root of the tree and the locations and effects of each mutation on the tree, the phenotypes at the tips are determined by adding mutant effects from the root to tip. To specify the root, we can assume without loss of generality that the ancestral phenotype for the entire population has a value 0 (this is similar to the common assumption in quantitative genetics literature that the ancestral state at each locus can be assigned a value of 0).

**Figure 1.**
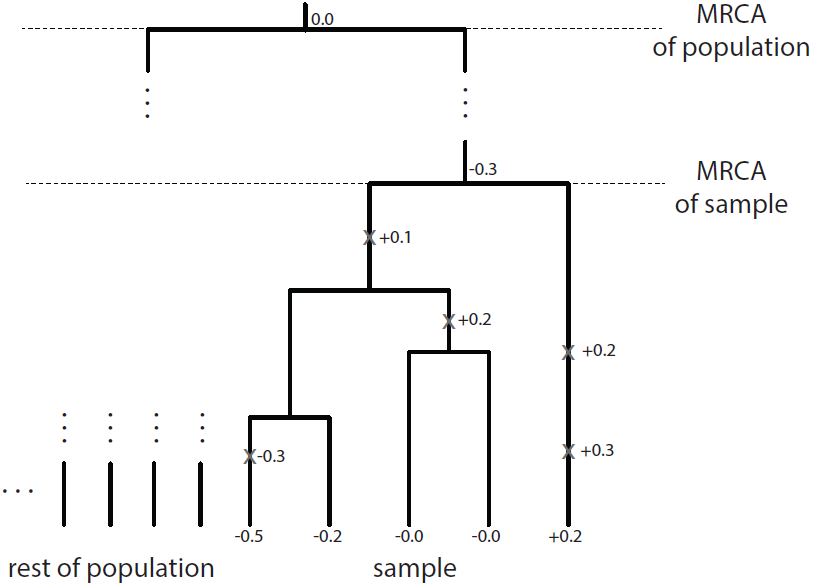
Example realization of coalescent process for a sample of size 5. Mutations (marked as light gray X’s), are placed upon the genealogy representing each individual in the population. Effects of each mutation are drawn from a probability distribution and are added along each branch length. The model is specified such that the most recent common ancestor (MRCA) of the population has phenotype 0.0, while the MRCA of the population may have a phenotype different from zero, due to mutations that accumulate between the MRCA of the sample and the MRCA of the population.

This mutational process can be described as a compound Poisson process (see also Khaitovich et al. (2005b); Chaix et al. (2008); Landis et al. (2013) for compound Poisson processes in a phylogenetic context). To ensure that this paper is self contained, we briefly review relevant facts about compound Poisson processes in the Appendix.

In the following, we ignore the impact of non-genetic variation and focus on the breeding value of individuals, i.e. the average phenotype of an individual harboring a given set of mutations.

## 3. Results

### 3.1. Computing the characteristic function of a sample

In many analyses, the object of interest is the joint probability of the data. If we let **X** = (*X*_1_, *X*_2_, …, *X_n_*) be the vector representing the quantitative traits observed in a sample of *n* individuals, we denote the joint probability of the data as *p*(*x*_1_, *x*_2_, …, *x_n_*). Note that, in general, *X_i_* and *X_j_* are correlated due to shared ancestry, and that *p* must be computed by integrating over all mutational histories consistent with the data. Hence, computing *p* directly is extremely difficult.

Instead, we compute the characteristic function of **X**. For a one-dimensional random variable, *X*, the characteristic function is defined as 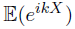 where *i* is the imaginary unit, *k* is a dummy variable. Generalizing this definition to an *n*-dimensional random variable, we are interested in computing

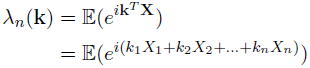

where **k** = (*k*_1_, *k*_2_, …, *k_n_*) is a vector of dummy variables. Like a probability density function, the characteristic function of **X** contains all the information about the distribution of **X**. Moreover, computing moments of **X** is reduced to calculating derivatives of the characteristic function, which will prove useful in the following.

We calculate this formula in two parts. First, we compute a recursive formula for *ϕ_n_*, the characteristic function given that ancestral phenotype of the *sample* is equal to 0. Then, we compute *ρ_n_*, the characteristic function of the ancestral phenotype of the sample, assuming that the characteristic function of the *population* is equal to 0. As we show in the Appendix, we can then multiply these characteristic functions to obtain the characteristic function of **X**.

We use a backward-forward argument to compute the recursive formula, first conditioning on the state when the first pair of lineages coalesce (backward in time) and then integrating (forward in time) to obtain the characteristic function for a sample of size *n*, *ϕ_n_*. This results in

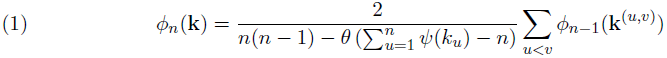

where **k**^(^*^u,v^*^)^ is the vector of length *n* − 1 made by removing *k_u_* and *k_v_* and adding *k_u_* + *k_v_* to the vector of dummy variables.

This equation has a straight-forward interpretation. The characteristic function for a sample of size *n*, *ϕ_n_*, is simply the characteristic function for a sample of size *n* − 1, *ϕ_n-_*_1_, averaged over all possible pairs that could coalesce first, multiplied by the characteristic function for the amount of trait change that occurs more recently than the first coalescent. The multiplication comes from the fact that the characteristic function of a sum of independent random variables is the product of the characteristic functions of those random variables. We prove this result in the Appendix (Section 5.3).

In the Appendix (Section 5.4), we also show that the characteristic function for the phenotype at the root of the sample is

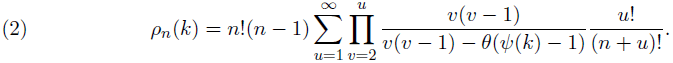

Intuitively, this equation arises by conditioning on whether *u* lineages are left in the *population* when the sample reaches its common ancestor and then averaging over the (random) time between when the individuals in the sample coalesce and when everyone in the population coalesces.

Hence, the characteristic function for a sample of size *n* is

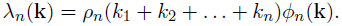

### 3.2. Sampling traits controlled by a small number of loci

It is common practice in both theoretical and applied quantitative genetics to summarize information about the phenotypic distribution within a population by computing central moments. However, care must be taken when interpreting theoretical predictions about central moments estimated from a sample. This is because the phenotypes in the sample are not independent, but instead correlated due to their shared genealogical history. Hence, in any *particular* population, an estimate of a central moment may deviate from its expected value, even as the number of individuals sampled grows to infinity (Aldous, 1985).

With this caveat in mind, we computed the first four expected central moments for a sample of phenotypes taken from this model (see Appendix for details). They are

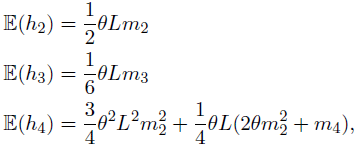

where *h_k_* is the unique minimum variance unbiased estimator of the *k*th central moment, *m_k_* is the *k*th moment of the mutational effect distribution and *L* is the number of loci that influence the trait.

These equations reveal that it may be possible to construct method-of-moments estimators for the moments of the mutation effect distribution and/or the number of loci that govern a trait.

### 3.3. “Infinitesimal” limits for large numbers of loci

Many traits are assumed to be governed by a large number of loci, each individually of small effect. This is known as an infinitesimal model (Falconer and Mackay, 1996). Typically, the sampling distribution in the infinitesimal limit is assumed to be Gaussian, by appealing to the central limit theorem. Here, we find that under certain circumstances traits may not be normally distributed, even in the limit.

To obtain a non-trivial limit, we must assume that as the number of loci controlling the trait increases, the effect of each individual locus decreases. Then, computing the characteristic function for a trait governed by a large number of independent loci is simple due to the fact the characteristic function of the sum of independent random variables is the product of their characteristic functions. Thus, assuming that each locus has the same effect distribution (this assumption can be relaxed relatively easily) the characteristic function of the limit distribution is given by

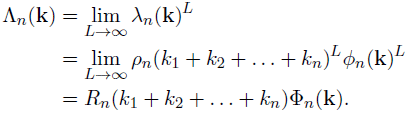

In the Appendix, we show that mutation effect distributions with power law behavior instead converge to a limiting *stable* distribution. A random variable *X* is said to have a power law distribution if *P* (*X > x*) ~ *κx^−α^* for large *x*, some *κ >* 0 and some *α* ∈ [0, 2). In this limit, individuals with shared genealogy may still have highly correlated phenotypes, due to rare mutations of large effect.

On the other hand, all mutation effect distributions without power law behavior converge to a Gaussian limit, due to the central limit theorem. In the Appendix, we show that samples taken from a population in this limit can be represented as a sample from a normal distribution with a random mean. In particular,

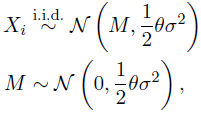

where 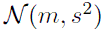 represents a normal distribution with mean *m* and variance *s*^2^.

### 3.4. Simulation

We simulated data to verify our analytical results and obtain some insight into the nature of the stable limiting distribution that arises for power law mutational effects. We first wanted to confirm that the trait distributions converge to univariate Gaussian limiting distributions as *n* → ∞ and *L* → ∞ when mutational kernels are not fat-tailed. To explore how the moments of the sampling distribution change with respect to *n*, *L*, and the mutational kernel, we asked for which values of *L* do the moments of the various mutational kernels leave a signature in the sampled quantitative traits. Finally, we conjectured that fat-tailed mutation kernels result in trait distributions that remain multimodal as *L* → ∞, which we verified by simulation rather than by mathematical proof.

For these simulation studies, we selected four mutational kernels: (1) the symmetric normal distribution for it’s simplicity, (2) the Laplace distribution because it is heavier-tailed (or more leptokurtic) than the normal distribution yet has finite variance, (3) the skew-normal distribution for it’s skewness parameter and tractability, and (4) the symmetric *α*-stable distribution because of its power-law behavior. To ensure that simulations of different non-fat-tailed distributions were comparable, we set the variance per locus to be *τ* ^2^ = *σ*^2^/*L* when we simulated *L* loci, meaning the trait distribution would have constant variance *θσ*^2^/2. Note the symmetric normal distribution is a special case of both the skew normal distribution when the skewness parameter is zero and the *α*-stable distribution when the “fat-tailedness” parameter is *α* = 2.

For all simulations, we generated coalescent genealogies and mutations using the program `ms` (Hudson, 2002). We then generated and mapped mutational effects using custom scripts in `R` (R Core Team, 2013).

Code is available at http://github.com/Schraiber/quant trait coalescent.

#### 3.4.1. Univariate Gaussian limit

For mutational kernels of small effect size and variance *τ*^2^ per locus, the sampling distribution converges to a normal distribution with variance *θσ*^2^/2 where *Lτ*^2^ → *σ*^2^ as *L* → ∞. We simulated 100 replicates of trait data for *L* ∈ {1, 2, 4, …, 256} and *n* ∈ {2, 4, 8 …, 512} with mutation parameters *θ* = 2 and *τ* = 1 for the normal, the skew-normal (skewness = 0.9), and the Laplace distributions. We then assessed convergence to the normal limit using the Kolmogorov-Smirnov (KS) test statistic, *D*, which equals zero when two distributions are identical. Figure 2 reports the frequency we reject the null hypothesis—that the limiting and sampled distributions are identical—for each batch of 100 replicates per value of *n* and *L* for *p*-values less than 0.05. For *n* ≤ 4, the KS test lacked power to reject the null hypothesis whatsoever. For *n* ≥ 8, the three mutational kernels converge to the limiting normal distribution in a similar fashion, with the sampling and limiting distributions bearing strong resemblance when *L* > 16. Distinctly, the Laplace distribution converges to normality at a slower rate than other mutational kernels, likely resulting from it being leptokurtic (Figure 3).

**Figure 2.**
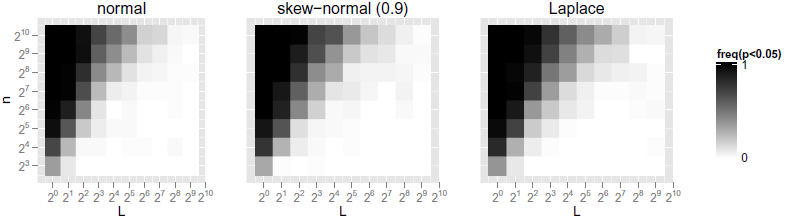
Frequency to reject normal sampling distribution. Heatmap cells correspond to number of sampled individuals, *n*, and number of loci, *L*. Panels are labeled with their respective mutational kernels. For 100 simulated replicates per cell, heatmap values correspond to the frequency the Kolmogorov-Smirnov test rejects the null hypothesis (*p* < 0.05) that the sampling distribution and the limiting normal distribution are equal. White cells indicate the sampling distribution looks normal. Black cells indicate the sampling distribution does not look normal.

**Figure 3.**
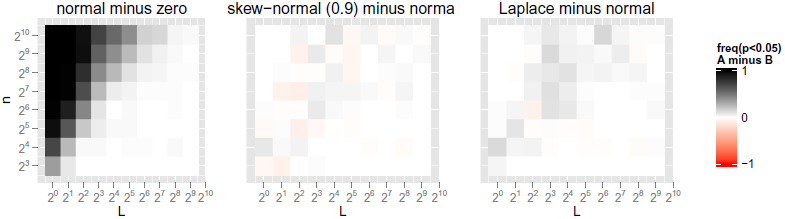
Comparison of frequency to reject normal sampling distribution. Heatmap cells correspond to number of sampled individuals, *n*, and number of loci, *L*. Panels are labeled with their respective mutational kernels. For 100 simulated replicates per cell, heatmap values correspond to the frequency the Kolmogorov-Smirnov test rejects the null hypothesis (*p <* 0.05) that the sampling distribution, *A*, and the limiting normal distribution are equal *minus* the frequencies computed for a second distribution, *B*. White cells indicate both distributions report equal frequencies. Black cells indicate *A* looks normal more often than *B*. Red cells indicate *A* looks normal less often than *B*.

#### 3.4.2. Central moments

We assessed the signature left by various mutational kernels on the sampling distribution by computing the central moments across simulation replicates. While the variance (*h*_2_) remains constant for all values of *L* regardless of the mutational kernel (by experimental design), the third central moment (*h*_3_) and the fourth central moment (*h*_4_) depend on the mutational kernel for small values of *L*. As *L* → ∞, the sample moments converge to those of a normal distribution. Here, we characterize the deviation from the normally-distributed moments under a variety of mutational kernels: the (symmetric) normal distribution; the skew-normal distribution for skewnesses 0.1, 0.5 and 0.9; and the Laplace distribution. We omitted the *α*-stable distribution from this portion of the study since its moments *h*_2_, *h*_3_, and *h*_4_ only exist when *α* = 2, i.e. when it is Gaussian.

We simulated data while varying the number of loci, *L* ∈ {1, 2, 4, …, 256}, holding the sample size constant, *n* = 1024, for 2000 replicates for each of the five mutation kernels. Afterwards, we computed the mean *h*_2_, *h*_3_, and *h*_4_ statistics across replicates of each mutation kernel and value of *L* for comparison with their expected *h*-statistic values (Figure 4). As expected, *h*_2_ remains constant regardless of the mutation kernel or *L*. The normal and Laplace distributions are symmetric and produce sample *h*_3_ values near zero, indicating no skewness. The skew-normal mutation kernel result in non-zero skewness even for traits controlled by over 100 loci so long as the kernel is sufficiently strongly skewed. The speed the sampling distribution’s third central moment, *h*_3_, converges to zero in inverse proportion to the magnitude of its mutational kernel’s skewness value. All distributions produce non-zero *h*_4_ values when *L* is small, due to the randomness of the mutation process. The *h*_4_ value of the Laplace distribution, the sole leptokurtic mutational kernel in this comparison, is the slowest of all kernels to converge to the normal limit.

**Figure 4.**
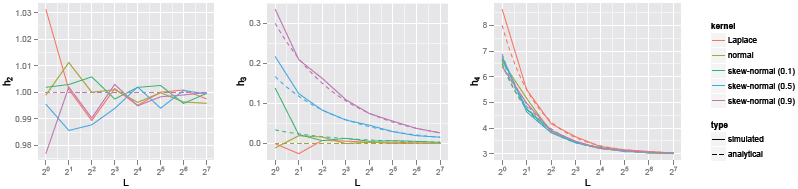
Central moments. From left to right, the panels correspond to the central moments, *h*_2_, *h*_3_, and *h*_4_, respectively, for the sampling distributions evolving under various mutational kernels. Data were simulated for 1024 sampled individuals and 2000 replicates for eight values of *L*, the number of loci. Colors distinguish the mutational kernel and relevant kernel parameters (if any). Solid lines correspond to moment values computed from the simulated data. Dashed lines correspond to the expected moment values.

#### 3.4.3. Multimodality

As *n* → ∞ and *L* → ∞, we proved that sampling distributions generated by finite-variance mutational kernels converge to the unimodal normal distribution and conjectured that power-law mutational kernels, such as the *α*-stable, converge to multimodal stable distributions. Here, we demonstrate by simulation our proven and conjectured modality results hold as *L* → ∞.

To do so, we test for unimodality using the dip statistic, *D* (Hartigan and Hartigan, 1985). Briefly, *D*(*F*_0_, *F*_1_) gives the minimized maximum difference between an empirical distribution, *F*_1_, and some unimodal (null) distribution, *F*_0_, where *F*_0_ is typically taken to be the uniform distribution. *D* approaches zero when *F*_1_ is unimodal and equals 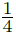 when the distribution is perfectly bimodal (i.e. two point masses). We used the R package `diptest` (Maechler and Ringach, 2012) to compute *p*-values for each simulated dataset, recording the frequency of replicates whose *p*-value is less than 0.05 for each mutational kernel and value of *L*. If the limiting distribution is unimodal, we expect this frequency to be less than 0.05 as *L* increases. Conversely, we expect multimodal limiting distributions to converge in frequency to some value greater than 0.05 as *L* increases.

We complemented the simulated data from Section 3.4.2 with three additional *α*-stable mutational kernels for *α* ∈ {1.5, 1.7, 1.9}, keeping the coalescent-mutation process variance equal across all datasets.

Figure 5 shows the trait distributions under *α*-stable mutation kernels remained multimodal as *L* → ∞ and stratify according to their respective *α* values: as *α* decreases the large-effect mutations responsible for multimodality grow more prominent. Mutation kernels of small effect size become unimodal as *L* → ∞. Notably, Laplace-distributed mutations converge to unimodality more slowly the normally-distributed mutations, echoing the results reported in Sections 3.4.1 and 3.4.2. Also note that when the number of loci is small (*L* ≤ 4) the sampling distribution is multimodal regardless of the mutation kernel. This corroborates our earlier KS tests (Section 3.4.1), which found simulated data for *L* ≤ 4 bore little to no resemblance to a unimodal normal distribution.

**Figure 5.**
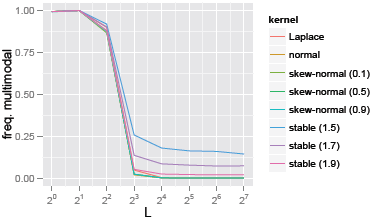
Frequency to reject unimodal sampling distribution. Solid lines report the frequency the null hypothesis of the dip-test, that the sampling distribution was unimodal, was rejected for *p* < 0.05 when evolving under various mutational kernels. Data were simulated for 1024 sampled individuals and 2000 replicates for eight values of *L*, the number of loci. Colors distinguish the mutational kernel and relevant kernel parameters (if any).

### 3.5. *Neurospora crassa* gene expression

Our simulations show that skewness, leptokurtosis, and multimodality may surface in the sampling distribution of quantitative traits, so we searched for these patterns in the *N. crassa* gene expression data reported by Ellison et al. (2011). Based on our modeling framework, the details of the deviation from normality can be used to infer the characteristics of the underlying mutational kernel. We selected this dataset because the data were collected so as to minimize environmental effects, because many changes gene expression may only be weakly deleterious if not neutral, and because transcriptomes contain thousands of comparable and consistently measurable quantitative traits. For these analyses, we only look at properties of the sampling distribution and make no assumptions about the generating process.

Forty-eight individuals were sampled from a wild population in Louisiana. The samples were then propagated in a controlled laboratory setting to minimize environmental and genotype-by-environmental effects on the quantitative traits. RNAseq raw read counts were obtained for 9793 genes, then normalized using upper quartile normalization (raw read counts divided by transcript length, further divided by the third quartile of all ranked read counts per individual). All expression levels were log-transformed. To control for noise, we discarded any weakly expressed gene with values less than log(-5.0) for any of the 48 individuals, then additionally discarded the most extreme value per gene, yielding 7079 genes with 47 individuals per gene. We expect these noise-control filters to bias our trait distribution toward normality, in particular, towards unimodal symmetric distributions with no excess kurtosis.

We used the Shapiro-Wilk test to assess normality and the dip test to assess multimodality. Additionally, we computed the sample skewness and kurtosis. Twenty-five and 697 genes reported p-values less than 0.05 for the dip test and Shapiro-Wilk tests, respectively, 14 of which fell into both categories (Figure 6A). We saw that genes in which normality is rejected tend to be positively and negatively skewed with approximately the same frequency (Figure 6B), and are more often leptokurtic than platykurtic (Figure 6C) For genes where we failed to reject normality, the mean sample skewness is −0.016 (i.e. mildly negatively skewed) and mean sample kurtosis is 2.78 (i.e. mildly platykurtic). After correcting for a false discovery rate of 10%, we identified one multimodal trait and 116 non-normally distributed traits. Among these discoveries, 57 were negatively skewed while 59 were positively skewed, and 23 were platykurtic while 93 were leptokurtic. Figure 7 shows the sampled trait distributions for six of the 116 non-normally distributed genes. The complete list of 116 genes is available at https://github.com/Schraiber/quant_trait_coalescent.

**Figure 6.**
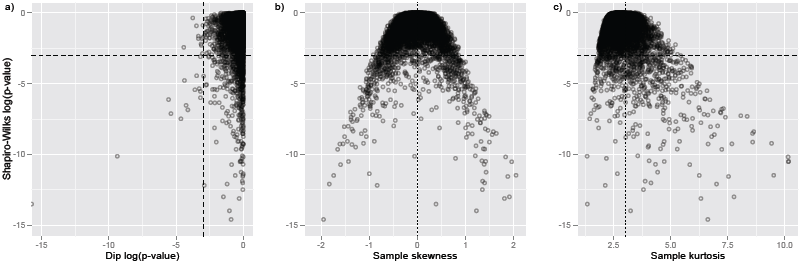
*N. crassa* gene expression trait distributions. All panels share a common y-axis, the log of the Shapiro-Wilk test’s p-value, where values less than log(0.05) are below the dashed horizontal line, indicating the rejection of normality. A) The log of the dip test’s p-value, where values less than log(0.05) are to the left of the dashed vertical line, meaning the rejection of unimodality. B) Skewness, where the dotted line shows the expected skewness under normality, 0. C) Kurtosis, where the dotted line shows the expected kurtosis under normality, 3.

**Figure 7.**
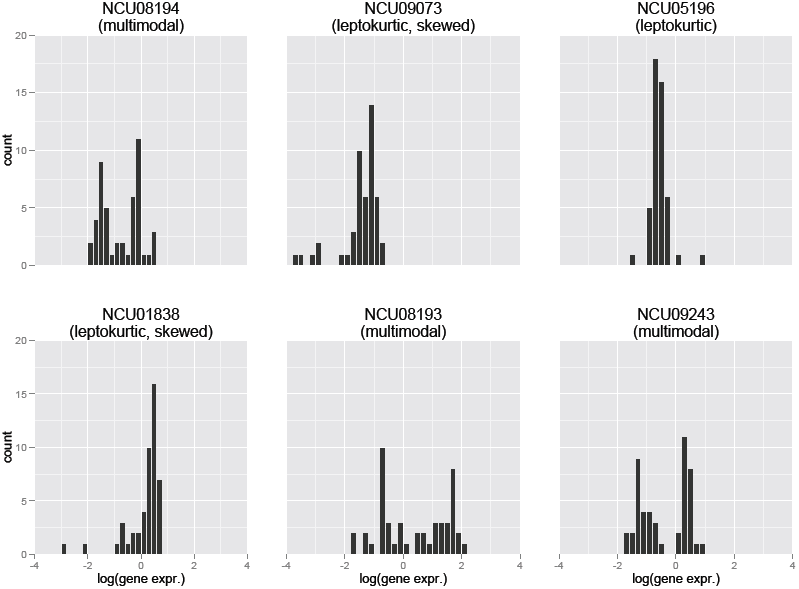
Examples of non-normal gene expression trait distributions. One hundred sixteen gene expression trait distributions (of 7079 genes) were found to be significantly non-normal under the Shapiro-Wilk test with a false discovery rate of *α* = 0.1. Of the 116 genes, six genes (NCU08194, NCU09073, NCU05196, NCU01838, NCU08193, NCU09243) that strongly rejected normality were chosen to represent the variety of skewed, leptokurtic, and multimodal distributions sampled for many genes.

The Shapiro-Wilk and dip tests may have insufficient power to reliably reject normality and unimodality hypotheses when given only 48 samples (49 samples minus the one most extreme-valued sample). We assessed power by simulation in a few simple cases. For example, for 48 samples drawn from an *α*-stable with *α* = 1.8, approximately 41% rejected normality under the Shapiro-Wilk test. Similarly, for a skew-normal distribution with skewness 0.5, 23% rejected normality. A simple bimodial distribution with equal mixture weights, means (0, 4), and standard deviations (1, 1) rejected the dip test 43% of the time. When one mixture component outweighs the second component four-fold (i.e. when a minor clade in the population carries a mutation of large effect), the dip-test is rejected only 0.71% of the time.

## Discussion

The natural world is replete with quantitative trait variation and understanding the forces governing their evolution is a central goal of evolutionary biology. The model of Fisher (1918), which explained how quantitative variation can be generated by Mendelian inheritance, provides an underpinning for understanding the generation and maintenance of variation in continuous characters. A primary assumption of much of this work is traits are controlled by a large number of loci and that new mutations have a very small, symmetric effect on the trait value.

In this work, we introduced a coalescent framework for modeling neutral evolution in quantitative traits. This stands in contrast to past work, which has typically taken a forward-in-time approach based on classical population genetics (but see Whitlock (1999) who also utilized a coalescent model). Our backward-in-time, sample-focused approach enabled us to derived an expression for the joint distribution of the data with arbitrary mutational effects and numbers of loci. We found that traits governed by a large number of loci with small effects are well-modeled by a Gaussian distribution, as expected. However, we saw that with small numbers of loci, significant departures from normality can be observed. Moreover, for fat-tailed (or power-law) mutational kernels, there are significant departures from normality (including multi-modality), even when the number of loci becomes large.

We assessed departure from normality in traits governed by a small number of loci by exploring the central moments of three different mutational kernels (normal, skew-normal and Laplace distributions) both analytically and by simulation. We showed that although all three mutational kernels converge to a Gaussian distribution, traits controlled by a small number of loci retain the signature of their underlying mutational kernel in their 3rd and 4th central moments. Hence, it may be possible to reconstruct aspects of the mutational effect distribution by observing phenotypes in natural populations. This may be particularly interesting for analyzing variation in gene expression, because mutational effects in *cis* may be strongly skewed (Khaitovich et al., 2005a; Chaix et al., 2008; Gruber et al., 2012). Our theory suggests that the distribution of gene expression in a population might therefore be skewed.

We were also interested in the circumstances under which multi-modal phenotypic distributions can arise. When a trait has a simple genetic architecture, it’s easy to see that there must be discrete phenotypic clusters, corresponding to groups of individuals sharing the same mutations. As the number of loci increases, there are more mutational targets (and thus more mutation events), which smooths the distribution, causing the sampling distribution to converge to the appropriate limiting distribution. For mutational effects with finite variance, this ultimately results in a limiting Gaussian distribution, consistent with the central limit theorem. However, when the mutational kernel is fat-tailed, the marginal effects of each locus do not vanish as the number of loci grows. Thus, some clade-specific mutations will always be of large effect despite the number of loci assumed by the model, resulting in a multi-modal sampling distribution.

Following our simulations, we asked whether the sampling distributions for empirical quantitative trait data were testably non-normal, as might be generated by the neutral model we presented earlier. For a sample of 49 strains of *N. crassa*, we found 116 of 7079 genes were detectably non-normal with a false discovery rate of 10%. Qualitatively, many more than 116 genes appeared non-normal, but more than 49 samples are needed for sufficient testing power. Because the data were generated while controlling for environmental effects then filtered for noisy measurements, we expect the quantitative trait variation must be explained predominantly by genetic factors. Several genes showed evidence of skewness, which our model shows could result from a skewed mutant effect distribution. Similarly, many genes showed evidence for leptokurtosis, although this may be due to either a simple genetic architecture (i.e. the trait is controlled by few loci) or a truly leptokurtic mutational kernel. The neutral model we proposed describes gene expression evolution only when no selection is acting on the quantitative trait, though we suspect that many of our results will hold qualitatively under weak selection. Of course, the assumption of weak or no selection is unlikely to be true across the entire transcriptome. Nonetheless, we believe that these qualitative aspects of the data can be used to shed light on the underlying mutational processes governing quantitative trait evolution. Whatever genetic process generated these data, e.g. an adaptive model, an adequate model must be capable of explaining skewed, leptokurtic, and multimodal sampling distributions.

These results show that even under the assumption of neutrality, significant departures from normality are possible and can be detected in empirical data. It is possible that these deviations from normality may be conflated with signatures of selection acting on quantitative variation. Several recent studies have claimed that evidence of non-Gaussianity may be evidence for non-neutral evolution at macroevolutionary time scales. For instance, Khaitovich et al. (2005a); Chaix et al. (2008) found that the distribution of gene expression differences between great apes is strongly positively skewed. Similarly, Uyeda et al. (2011) argued that there is a one million year wait between bursts of evolution in the fossil record and numerous studies have explored non-Gaussian trait divergence in a phylogenetic context (Landis et al., 2013; Eastman et al., 2013). While it is unlikely that the population genetic model we developed can be directly applied to macroevolutionary data of this sort (Estes and Arnold, 2007), it is important to recognize that such effects can be due to purely neutral processes.

On shorter time scales, there is significant interest in detecting non-neutral quantitative trait evolution among closely related species or populations. One powerful method compares a measure of quantitative trait divergence, *Q_st_*, to the fixation index, *F_st_* (McKay and Latta, 2002; Ovaskainen et al., 2011). However, this requires estimates of breeding values from common-garden experiments, and may be difficult to achieve. In other cases (e.g. Lemos et al. (2005)) more phenomenological approaches are taken, by comparing within and between species phenotypic diversity. The null distributions of these approaches typically rely on assumptions of the infinitesimal model, which we have shown may be violated due to mutations of large effect and/or loci with relatively simple genetic bases. To address these issues and leverage the abundance of modern quantitative trait data, Berg and Coop (2014) developed a method that explicitly uses breeding values estimated from quantitative trait mapping studies. When such effect size estimates are unavailable, it may be possible to use our formalism to develop robust null models to detect selection.

Our coalescent approach can be extended in several ways. Notably, we consider only haploid populations. In principle, an extension to diploid individuals is straight-forward using the result of Möhle (1998) that diploid, dioecious populations of size *N* are readily modeled by pairing random chromosomes from a haploid population of size 2*N*. To incorporate diploidy, we would also need to incorporate a model of dominance, of which several exist in the literature (e.g. the model of independent dominance of Lynch and Hill (1986).

From the point of view of the coalescent process, it is straightforward to apply our model to populations that have undergone complex demographic histories. This is because the dynamics of a coalescent under population size fluctuations and population structure are well known. Moreover, we explored only unlinked, neutral loci and it may be possible to obtain some analytical results for linked loci and/or weak natural selection by using the ancestral recombination graph and ancestral selection graph, respectively. While analytical results are difficult within these frameworks, we believe that they can be used to perform simple simulations of quantitative traits evolving in complex scenarios, thus enabling Approximate Bayesian Computation.

## 5. Appendix

### 5.1. Compound Poisson processes

To obtain the probability of the data under this model, we must be able to compute the probability of the change in phenotype along a branch of the tree. Unfortunately, except for very simple mutational models, this probability is impossible to compute analytically. Instead, we compute the characteristic function of the change along a branch.

Using standard results for compound Poisson processes (Kingman, 1992), we see that the characteristic function of the change along a branch of length *t* (in coalescent units) is

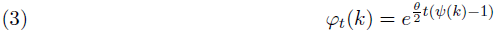

where *ψ* is the characteristic function of the mutational effect distribution.

### 5.2. The phenotype at the root of the sample genealogy and the subsequent evolution within the sample are subindepenent

Note that

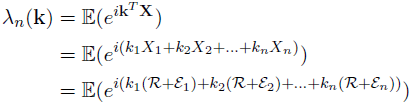

where 𝓡 is the phenotype at the root of the sample genealogy and *ε_u_* is the subsequent evolution leading to lineage *u* in the sample. So,

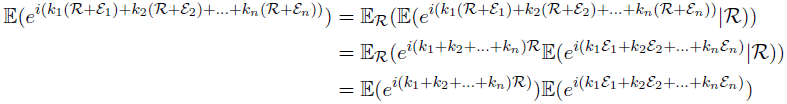

where the last line follows by independent and stationary increments of the compound Poisson process. Thus, 𝓡 and (*ε*_1_, *ε*_2_, …, *ε_n_*) subindependent, and hence their joint characteristic function is the product of their characteristic functions.

### 5.3. Proof of recursive formula for the characteristic function

First, we condition on the state at the first coalescence (going back in time). The state consists of three components: 1) which pair of individuals coalesce, (*u, v*), 2) the time of the coalescent event, *T_c_*, and 3) the trait value in each lineage at that time, **X**^′^ (note that, given (*u, v*), we have that *X_u_^′^* = *X_v_^′^*, since those two ilneages have coalesced and hence had the same trait value at the time of coalescence). Then,

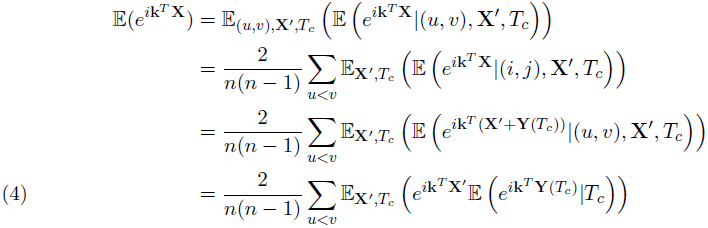

where **Y**(*t*) = (*Y*_1_(*t*), *Y*_2_(*t*), …, *Y_n_*(*t*)) is the vector accounting for the evolution on each lineage that occurs during time *t*. The second line follows by the fact that each pair is equally likely to coalesce (with probability 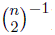) and the third line by independent increments of a compound Poisson process.

Now, we compute the internal expectation going forward in time. Noticing that 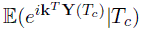 is simply the characteristic function of a compound Poisson process run for length *T_c_*, we see from (3) that

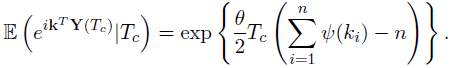

Because *T_c_* and **X***^′^* are independent, we can integrate over *T_c_* analytically in the outer expectation. The distribution of the time to the first coalescent event in a sample of size *n* is Exponential with rate 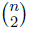, hence,

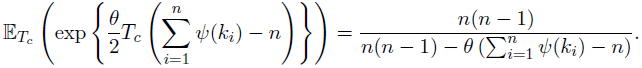

Plugging this result into (4) results in

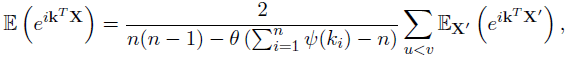

but since **X***^′^* is simply the result of the same process where two of the entries are identical, we obtain the recursive formula (1).

To initialize the recursion, we must compute the characteristic function for a sample of size 2. This is

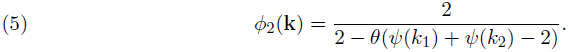

### 5.4. The phenotype at the root of the sample genealogy

First, we note that, conditional on the time between when the sample genealogy finds a common ancestral and the population genealogy finds a common ancestor, Δ, the characteristic function of the phenotype at the root of the sample genealogy is

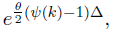

by using equation (3). Thus, the after integrating over Δ, the desired quantity is *moment generating function* of Δ, defined by

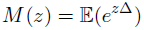

evaluated as 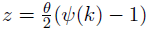.

We compute *M* (*z*) by conditioning on how many lineages are left in the population genealogy when the sample reaches its most recent common ancestor. To do this, we make use of a result of Saunders et al. (1984),

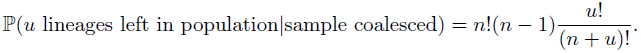

Given that *u* lineages are left in the population when the sample reaches its most recent common ancestor, the remaining time until the whole population reaches its common ancestor is simply the time it takes for a coalescent started with *u* to reach its most recent common ancestor, *C_u_*. Thus,

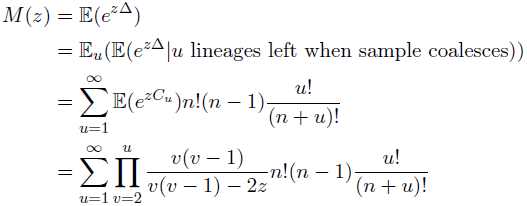

where the final line follows by recognizing that *C_u_* is the sum of *u* − 1 independent exponential random variables with means 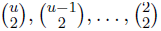. Substituting 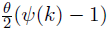 for *z* yields the desired result.

### 5.5. Computing sample central moments

While it is difficult to compute the expectation of any sample central moments for a particular sample, it is possible to average over replicate populations to compute expectations. This results in

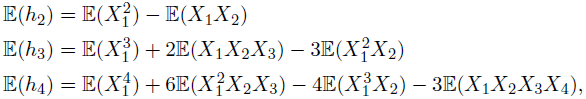

where the expectations on the right hand sides are over the *correlated* phenotypes in the sample. It is possible to compute these expectations by taking derivatives of the characteristic function (1). After simplifying, one then arrives at the formulas in the main text.

### 5.6. Derivation of multivariate stable limit for sample distribution

Recall that a random variable *X* is said to have a fat-tailed (or power-law) distribution if

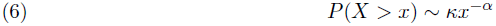

for large *x* and some *κ >* 0. As is typical in the literature, we reserve the term “fat-tailed” for distributions with *α* ∈ (0, 2).

To obtain an appropriate scaling limit, we assume that there is a parameter *t*, related to the parameter *κ* in (6) by

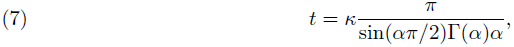

such that *Lt* → *s* as *n* → ∞. The parameter *s* is related to the scale parameter of the resulting limit distribution.

We provide a heuristic derivation, rather than a rigorous proof. First, we argue by induction that the (per locus) characteristic function for a sample of size *n* is

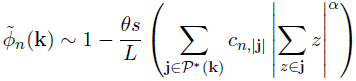

for large *L*, where 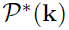 is the power set of the elements in **k**, *except* the set {*k*_1_, *k*_2_, …, **k*_n_*}, and *c_n_*|**j**| is a combinatorial constant that depends only on the sample size *n* and |**j**|, the size of the set **j**.

Note that for *n* = 2, this can be seen by observing that for large *L*, the characteristic function of a fat-tailed distribution is asymptotically

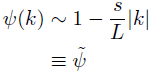

Thus,

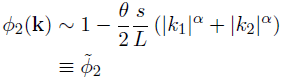

Now, assume that the formula holds for 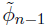. Using the recursion (1), we have

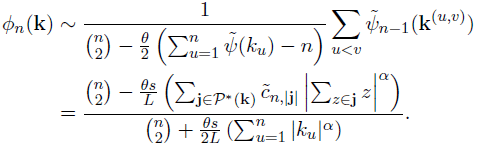

The second line follows from plugging 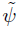 and 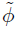, and 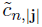 arises by summing over the appropriate terms coming from all characteristic functions in the sum. Again looking for an asymptotic for large *L*, we see that

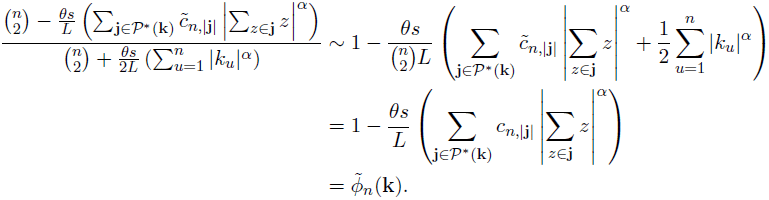

Finally, we note that by raising 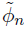 to the *L*th power, and taking the limit as *L* → ∞, we obtain the log characteristic function

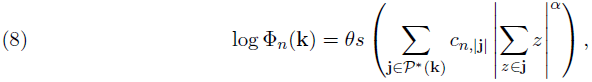

where all terms are defined as before.

The characteristic function in (8) can be recognized to be that of a multivariate *α*-stable distribution (Press, 1972). These multivariate distributions are fat-tailed generalizations of the familiar multivariate normal distribution, and this limit corresponds to a generalized multivariate central limit theorem for sums of random vectors with fat-tailed distributions.

### 5.7. Limiting distribution of the phenotype at the root of the sample genealogy

Again, we proceed heuristically rather than rigorously. First, note that for large *L*,

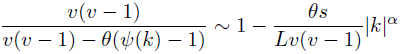

so that

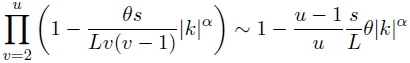

for large *L*. Thus,

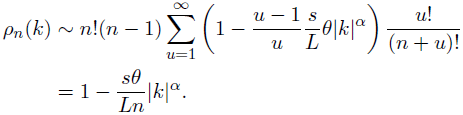

So by definition of the exponential function, we have that

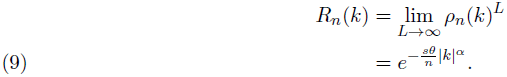

which the characteristic function of a univariate *α*-stable distribution, arising from the fact that the phenotype at the root of the sample genealogy is itself a limit of a sum of random variables. Note that as *n* → ∞ (i.e. the sample becomes the whole population), *R*(*k*) → 1, because the root of the sample genealogy is the same as the root of the population genealogy and the root value has been specified to be equal to 0.

### 5.8. Multivariate Gaussian limits

For the case where the mutation distribution is not fat-tailed, we can use the multivariate central limit theorem to more efficiently derive the limiting distribution. The appropriate scaling in this case is to assume that if *τ*^2^ is the variance of the mutation effect kernel, then *Lτ*^2^ → *σ*^2^ as *L* → ∞.

To apply the multivariate central limit theorem, we must derive the pairwise covariances between samples. While the required covariances could be computed by taking derivatives of the characteristic function, it is more instructive to compute these moments directly. For simplicity, we assume that the mutation effect distribution has mean 0 and variance *τ*^2^.

Assume that the population genealogy at a single locus, 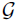, is fixed. Noting that the variance per unit time accrued by the mutational process is *θ/*2*τ*^2^ and using the rules for calculating covariance structure on a phylogeny, it’s easy to see that for samples *i* and *j* we have

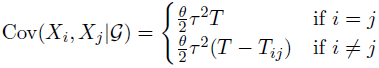

where *T* is the height of 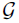 and and *T_ij_* is the height of the most recent common ancestor of samples *i* and *j*. We can then use the law of total covariance,

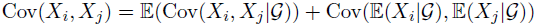

to see that

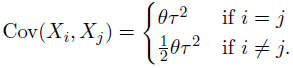

This arises because 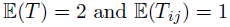.

Hence, as the number of loci increases to infinity in such a way that *Lτ*^2^ → *σ*^2^, the sampling distribution converges to a multivariate normal distribution with mean 0 and variance covariance matrix Σ having elements

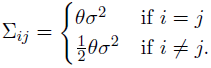

Because the pairwise covariances are equal, the random vector **X** is an exchangeable Gaussian random vector. Hence, using well-known facts about the representation of exchangeable Gaussian random vectors, one arrives at the representation in the main text.

## Acknowledgments

We are grateful to Monty Slatkin, Anand Bhaskar and Matt Pennell for reading an earlier version of this manuscript and providing extremely detailed suggestions that significantly improved its clarity. We are also indebted to Anand Bhaskar for suggesting the forward-backward approach that led to (1). We owe a debt of gratitude to Chris Ellison for his assistance with the *N. crassa* data. J.G.S. was supported by National Institutes of Health grants R01-GM40282 (awarded to Montgomery Slatkin) and National Science Foundational postdoctoral fellowship DBI-1402120. M.J.L. was supported by National Institutes of Health grant R01-GM069801 (awarded to John P. Huelsenbeck).

## References

Aldous, D. J. 1985. Exchangeability and Related Topics. Springer.

Berg, J. J. and G. Coop. 2014. A population genetic signal of polygenic adaptation. PLoS Genetics 10:e1004412.

Bürger, R. and R. Lande. 1994. On the distribution of the mean and variance of a quantitative trait under mutation-selection-drift balance. Genetics 138:901–912.

Chaix, R., M. Somel, D. P. Kreil, P. Khaitovich, and G. Lunter. 2008. Evolution of primate gene expression: drift and corrective sweeps? Genetics 180:1379–1389.

Chakraborty, R. and M. Nei. 1982. Genetic differentiation of quantitative characters between populations or species: I. Mutation and random genetic drift. Genetical Research 39:303–314.

Doebeli, M., H. J. Blok, O. Leimar, and U. Dieckmann. 2007. Multimodal pattern formation in phenotype distributions of sexual populations. Proceedings of the Royal Society B: Biological Sciences 274:347–357.

Eastman, J. M., D. Wegmann, C. Leuenberger, and L. J. Harmon. 2013. Simpsonian ‘evolution by jumps’ in an adaptive radiation of anolis lizards. arXiv preprint arXiv:1305.4216.

Ellison, C. E., C. Hall, D. Kowbel, J. Welch, R. B. Brem, N. Glass, and J. W. Taylor. 2011. Population genomics and local adaptation in wild isolates of a model microbial eukaryote. Proceedings of the National Academy of Sciences 108:2831–2836.

Estes, S. and S. J. Arnold. 2007. Resolving the paradox of stasis: models with stabilizing selection explain evolutionary divergence on all timescales. The American Naturalist 169:227–244.

Falconer, D. and T. Mackay. 1996. Introduction to Quantitative Genetics. 4 ed. American Genetic Association.

Fisher, R. 1930. The Genetical Theory of Natural Selection. Clarendon Press.

Fisher, R. A. 1918. The correlation between relatives on the supposition of Mendelian inheritance. Transactions of the Royal Society of Edinburgh 52:399–433.

Galton, F. 1883. Inquiries into Human Faculty and its Development. Macmillan.

Galton, F. 1889. Natural Inheritance. Macmillan.

Gruber, J. D., K. Vogel, G. Kalay, and P. J. Wittkopp. 2012. Contrasting properties of genespecific regulatory, coding, and copy number mutations in *Saccharomyces cerevisiae*: frequency, effects, and dominance. PLoS genetics 8:e1002497.

Haldane, J. 1954. The statics of evolution. Evolution as a Process Pages 109–121.

Hartigan, J. A. and P. Hartigan. 1985. The dip test of unimodality. The Annals of Statistics Pages 70–84.

Hudson, R. R. 2002. Generating samples under a Wright–Fisher neutral model of genetic variation. Bioinformatics 18:337–338.

Khaitovich, P., I. Hellmann, W. Enard, K. Nowick, M. Leinweber, H. Franz, G. Weiss, M. Lachmann, and S. Pääbo. 2005a. Parallel patterns of evolution in the genomes and transcriptomes of humans and chimpanzees. Science 309:1850–1854.

Khaitovich, P., S. Pääbo, and G. Weiss. 2005b. Toward a neutral evolutionary model of gene expression. Genetics 170:929–939.

Kimura, M. 1965. A stochastic model concerning the maintenance of genetic variability in quantitative characters. Proceedings of the National Academy of Sciences of the United States of America 54:731.

Kingman, J. F. C. 1992. Poisson Processes. 3 ed. Oxford University Press.

Kopp, M. and J. Hermisson. 2006. The evolution of genetic architecture under frequency-dependent disruptive selection. Evolution 60:1537–1550.

Lande, R. 1976. Natural selection and random genetic drift in phenotypic evolution. Evolution Pages 314–334.

Landis, M. J., J. G. Schraiber, and M. Liang. 2013. Phylogenetic analysis using Lévy processes: finding jumps in the evolution of continuous traits. Systematic biology 62:193– 204.

Latter, B. 1960. Natural selection for an intermediate optimum. Australian Journal of Biological Sciences 13:30–35.

Latter, B. 1970. Selection in finite populations with multiple alleles. ii. Centripetal selection, mutation, and isoallelic variation. Genetics 66:165.

Lemos, B., C. D. Meiklejohn, M. Cáceres, and D. L. Hartl. 2005. Rates of divergence in gene expression profiles of primates, mice, and flies: stabilizing selection and variability among functional categories. Evolution 59:126–137.

Lynch, M. and W. G. Hill. 1986. Phenotypic evolution by neutral mutation. Evolution Pages 915–935.

Mackay, T., R. F. Lyman, and M. S. Jackson. 1992. Effects of P element insertions on quantitative traits in *Drosophila melanogaster*. Genetics 130:315–332.

Maechler, M. and D. Ringach. 2012. diptest: Hartigans dip test statistic for unimodality.

McKay, J. K. and R. G. Latta. 2002. Adaptive population divergence: markers, qtl and traits. Trends in Ecology & Evolution 17:285–291.

Mendel, G. 1866. Versuche über pflanzenhybriden. Verhandlungen des naturforschenden Vereines in Brunn 4: 3 44.

Möhle, M. 1998. Coalescent results for two-sex population models. Advances in Applied Probability Pages 513–520.

Ovaskainen, O., M. Karhunen, C. Zheng, J. M. C. Arias, and J. Merilä. 2011. A new method to uncover signatures of divergent and stabilizing selection in quantitative traits. Genetics 189:621–632.

Pearson, K. 1894. Contributions to the mathematical theory of evolution. Philosophical Transactions of the Royal Society of London A. Pages 71–110.

Pearson, K. 1895. Contributions to the mathematical theory of evolution. III. Regression, heredity, and panmixia. Proceedings of the Royal Society of London 59:69–71.

Pearson, K. 1904. Mathematical contributions to the theory of evolution. XII. On a generalised theory of alternative inheritance, with special reference to Mendel’s laws. Philosophical Transactions of the Royal Society of London A. Pages 53–86.

Press, S. J. 1972. Multivariate stable distributions. Journal of Multivariate Analysis 2:444– 462.

R Core Team. 2013. R: A Language and Environment for Statistical Computing. R Foundation for Statistical Computing Vienna, Austria.

Saunders, I. W., S. Tavaré, and G. Watterson. 1984. On the genealogy of nested subsamples from a haploid population. Advances in Applied probability Pages 471–491.

Uyeda, J. C., T. F. Hansen, S. J. Arnold, and J. Pienaar. 2011. The million-year wait for macroevolutionary bursts. Proceedings of the National Academy of Sciences 108:15908– 15913.

Weldon, W. F. R. 1902. Mendel’s laws of alternative inheritance in peas. Biometrika Pages 228–254.

Whitlock, M. C. 1999. Neutral additive genetic variance in a metapopulation. Genetical research 74:215–221.

